# Quinomycin A reduces cyst progression in Polycystic Kidney Disease

**DOI:** 10.1101/2020.10.18.344689

**Authors:** Priyanka S Radadiya, Mackenzie M Thornton, Brenda Magenheimer, Dharmalingam Subramaniam, Pamela V Tran, James P Calvet, Darren P Wallace, Madhulika Sharma

**Author notes:** These authors contributed equally to the study (Priyanka S Radadiya, Mackenzie M Thornton). **Correspondence**, Madhulika Sharma, PhD, Department of Internal Medicine, University of Kansas Medical Center, Kansas City, KS 66160.

## Abstract

Polycystic kidney disease (PKD) is a genetic disorder that affects cilia homeostasis and causes progressive growth of tubular-derived cysts within the kidney. Efforts to find safer drugs for PKD have increased in the past few years after the successful launch of tolvaptan, the first approved drug to combat autosomal dominant PKD progression. Here we investigate the effects of Quinomycin A on progression of PKD. Quinomycin A is a bis-intercalator peptide that has previously shown to be effective against cancer progression. Quinomycin A treatment decreased cyst progression of human ADPKD primary renal epithelial cells grown in a 3D collagen gel to form cysts. In an orthologous mouse model of PKD, Quinomycin A administration reduced kidney to body weight ratios, and reduced cystogenesis. This was accompanied by decreased cell proliferation and fibrosis. Quinomycin treatments efficiently reduced the expression of Notch pathway proteins, RBPjk and HeyL in kidneys of PKD mice. Interestingly, Quinomycin treatments also normalized cilia lengths of collecting duct cyst-lining renal epithelia of PKD mice. This is the first preclinical study to our knowledge that demonstrates Quinomycin A has protective effects against PKD progression, in part by reducing Notch signaling and renal epithelial cilia lengths. Our findings suggest Quinomycin A has potential therapeutic value for PKD patients.

## Introduction

Polycystic kidney disease (PKD) is a genetic disease characterized by formation and expansion of renal cysts resulting from hyper proliferation and abnormal fluid secretion of renal epithelial cells (1, 2). Two common inherited forms of PKD are: **1)** Autosomal dominant polycystic kidney disease (ADPKD), a disease caused by mutations in either *PKD1* or *PKD2*, which encode polycystin1 (PC1) and polycystin 2 (PC2), respectively, and **2)** Autosomal recessive polycystic kidney disease (ARPKD), caused by mutations of *PKHD1*, which encodes fibrocystin. PC1, PC2 and fibrocystin form protein complexes that localize to the primary cilium, a non-motile sensory organelle, that transduces mechanical and/or chemical stimuli and mediates cellular signaling (3). Several signaling pathways are activated in response to mutations in PC1 and PC2 in ADPKD and efforts are devoted to target these pathways. Tolvaptan is presently the first line of treatment approved for ADPKD (4). Tolvaptan can inhibit cell proliferation, Cl-secretion, and cyst growth in renal cystic cells (ADPKD cells) stimulated by vasopressin (5). While the long-term safety of tolvaptan in patients with ADPKD is not clear, several alternative repurposed drugs are in clinical trials being tested for PKD; these include metformin, a drug for diabetes which acts by inhibiting glycolysis (6); niacinamide (a form of Vitamin B) has shown to inhibit sirtuin1 and reduces cyst growth in PKD mice (7) and tesevatinib, a tyrosine kinase inhibitor which inhibits epidermal growth factor (EGFR) and cyst growth (7, 8).

We have previously reported activation of Notch3 pathway in cystic epithelial cells of various mouse models of PKD as well as in human ADPKD patients (9). The Notch signaling pathway is activated upon binding of a Notch ligand to a Notch receptor, resulting in the gamma secretase-mediated release of the Notch intracellular domain (NICD). The NICD translocates to the nucleus and binds to recombination signal binding protein immunoglobulin kappa J region (RBPJk), which results in transcriptional activation of Notch target genes, *Hes* and *Hey*. Gamma secretase inhibitors (GSI) can inhibit Notch activity *in vivo* and *in vitro*, however these inhibitors have been shown to be associated with gastrointestinal toxicity in clinical trials (10). Thus, we investigated whether repurposing a drug, known to have inhibitory effects on Notch signaling, would attenuate PKD.

Quinomycin A, also called echinomycin is a member of quinoxaline family of antibiotics, originally isolated from Streptomyces echinatus (11). Quinomycin A can bind strongly to double stranded DNA and exhibits antitumor activities. Quinomycin treatments reduced acute myeloid leukemia in mouse model without any adverse effects on normal hematopoetic cells (12, 13). Quinomycin A also reduced disease progression in a mouse model of pancreatic cancer by targeting the Notch signaling pathway (14). Thus, we asked whether quinomycin can protect against PKD by inhibiting the Notch pathway activation.

We show that Quinomycin A can reduce cyst growth *in vitro* and *in vivo* and inhibits cell proliferation and fibrosis. Quinomycin A also inhibits aberrant activation of the Notch pathway, and interestingly, reduced cilia lengths of cyst-lining epithelial cells in an ADPKD mouse model. In contrast, Quinomycin A showed minimal effects on normal human renal epithelial cells and kidneys of control mice. Taken together, our data demonstrate that Quinomycin A selectively targets PKD cystic epithelial cells.

## Methods

### Antibodies and reagents

Antibodies and their sources are listed: Notch3 and HIF-1α from Abcam (Cambridge, MA), smooth muscle actin from (Sigma Aldrich, Saint Louis, MO), RBPjk (Santa Cruz, Dallas, TX), Hey L (Life Span Biosciences, Seattle, WA). Ki67 from Dako, Quinomycin A and anti-acetylated a tubulin from Sigma Aldrich (Saint Louis, MO). *Dolichus biflorus agglutinin* (DBA) was obtained from Vector Labs.

### Animal care and protocol

*Pkd1^RC/RC^: Pkd2^+/-^* (PKD) mice were generated by breeding *Pkd1^RC/RC^* mice with *Pkd2^+/-^* mice (both in C57BL/6J background) kindly provided by Drs. Peter Harris and Steven Somlo, respectively (15, 16). Homozygous *Pkd1^RC/RC^* and heterozygous *Pkd2^+/-^* mice have a slow progressing cyst phenotype. However, when these mice are bred to obtain *PKD1^RC/RC^, PKD2^+/-^* mice, cyst formation is accelerated. The *PKD1^RC/RC^, PKD2^+/-^* (PKD) mice are mildly cystic at birth and the cysts progress with age such that there is an exponential growth of the cysts between age P15 and P60 after which cyst growth is minimal (unpublished). We used these mice because the time period of cyst formation is moderate for the drug studies. Wildtype littermates without a PKD mutation *(Pkd1^+/+/^/Pkd2^+/+^)* were used as controls. Mice were generated at the PKD rodent core facility at the University of Kansas Medical Center.

### Quinomycin A treatments in mice

Quinomycin A was solubilized in distilled water and filter sterilized using a 0.2micron filter for treatments. Study groups consisted of (1) vehicle-treated wildtype (WT) mice. (2) Quinomycin A treated WT mice (3) vehicle treated PKD mice and (4) Quinomycin A treated PKD mice. Mice were weaned and treatments were started at postnatal day 22 for a total of 27 days with daily intraperitoneal injections of 10μg /kg body weight Quinomycin A. During euthanasia, mice were weighed and perfused with cold PBS after blood collection followed by collecting and weighing kidneys. One kidney was snap frozen and the other was fixed in 4% paraformaldehyde for 24hrs followed by storage in 70% ethanol at 4°C until blocking and sectioning for histology and immunohistochemistry.

### Histology, cystic Index and blood urea nitrogen (BUN) measurements

The fixed kidney tissues were processed at the core facilities of the University of Kansas Medical Center. Five-micrometer sections were stained with Hematoxylin and Eosin (H&E) as described previously (17). Cystic index was measured using ImageJ on H&E stained kidney sections. The area of each individual cyst within the section of entire kidney was calculated and then added together. This summed value was then divided by the total area of the section yielding in the value identified as the cystic index. This was done for a kidney section for each mouse used in the study in each PKD group. BUN was quantified using a Quantichrom Urea Assay Kit (Bioassay Systems), according to the manufacturer’s protocol.

### Human cell culture and treatments

Primary cultures of ADPKD and NHK epithelial cells were generated as described previously (18). Use of de-identified surgically discarded tissues complies with federal regulations and was approved by the Institutional Review Board at KUMC. ADPKD cells were obtained from multiple surface cysts ranging in size. NHK cells were cultured from sections of cortex. These cells have been shown to be enriched in collecting duct marker, *Dolichos biflorus agglutinin* (DBA) (19). Cells were cultured in DMEM/F-12 supplemented with 5% FBS, 5 μg/ml insulin, 5 μg/ml transferrin, and 5 ng/ml sodium selenite (ITS, Thermo Scientific) and penicillin (100 U/ml), streptomycin (130 μg/ml) (Pen/Strep)(20). Cultures were not passaged more than twice before being used in experiments. For treatments, ADPKD cells were grown to 70% confluency followed by a 24-hour low serum (0.001% and no ITS) treatment. Cells were then treated with vehicle or Quinomycin A for 4 days hours before studies.

### Cell viability assay

ADPKD or NHK cells were plated in a 12 well plate (20,000 cells/well) and allowed to grow to 70% confluency at 37° C under 5% CO2 followed by 24-hour low serum (0.001%) and no ITS treatment. The following day cells were treated with increasing concentration of Quinomycin A (0 to 10nM) . Cells were grown for 4 days and fresh media containing Quinomycin A was replaced every day. After 4 days, cells were trypsinized and pelleted. Cells were suspended in 500 μl media and cell viability tested in triplicates using cell counting kit-8 (CCK-8, Apex Biosciences, Houston, TX) and manufacturer’s instructions were followed. Viability was set at 100% for the vehicle control and relative values were calculated for other doses.

### *In vitro* 3D cyst assays

*In vitro* cyst assays were performed as described (21) (22, 23). Briefly, primary cultures of ADPKD cells were suspended in media containing type 1 collagen (PureCol, Advanced biomatrix, San Diego, CA) in a 96-well plate. Immediately after adding collagen, 100μl of media with collagen and cells (4× 10^3^ /100ml), was pipetted into each well. The plate was incubated at 37°C for 45 minutes to allow collagen to polymerize. Then, 150μl of defined media (1:1 DMEM/F12 with ITS, 5 × 10^−8^ M hydrocortisone, 5× 10^−5^ M triiodothyronine) containing 5 μM forskolin and 5 ng/ml EGF was added on to the polymerized gel to initiate cyst growth. Following formation of cysts at day 4, the agonists (FSK and EGF) were removed and the gels were rinsed twice with defined media. To initiate drug treatments, Quinomycin A at different concentrations was added to the wells with media. Fresh treatment media with drug was replaced every day for each treatment. After 4 days, the outer diameter of cross-sectional images of spherical cysts with distinct lumens were measured using a digital camera attached to an inverted microscope and analyzed with video analysis software (image pro-premier). Surface area was calculated from the outer diameters and total surface area of the cysts was determined from the sum of individual cysts within each well (6 wells per treatment). Cysts with diameters of 50μM or less were excluded. Data is presented as surface area/mm^2^.

### Immunohistochemistry (IHC)/ Immunofluorescence (IF)

IHC was performed as described previously (24). Briefly, kidney sections from wild type and PKD mice treated with Quinomycin A (quin) or vehicle (veh) were deparaffinized with xylene and hydrated with graded ethanol. Sections were then boiled in citrate buffer (10 mM sodium citrate, 0.05% tween 20, pH: 6.0) and cooled to room temperature. Sections were incubated for 30 min with 3% hydrogen peroxide for IHC or 0.5 M ammonium chloride for IF to block endogenous peroxidase or fluorescence activity, respectively. Subsequent washing in PBS and blocking with 10% normal serum (in PBS from the species the secondary antibody was raised in) for 1 h were followed by incubation for 1 h with primary antibodies in a humidified chamber. Slides were washed in PBS and incubated for 1 h in 1:400 diluted biotin-conjugated secondary antibodies (Vector Laboratories, Burlingame, CA) for IHC and 1:400 Alexa 488 or Alexa 594 (Invitrogen) for IF. Slides were washed in PBS again. For IF, the slides were cover slipped using prolong diamond (Invitrogen) and sealed with clear nail polish. For IHC, the slides were further incubated with avidin-biotin-peroxidase complex (ABC Elite; Vector Laboratories, Burlingame, CA) and detected with diaminobenzidine (DAB; Sigma Aldrich, St. Louis, MO). Tissue sections for IHC were then dehydrated with graded ethanol and mounted with permount before coverslipping (Fisher Scientific, Pittsburg, PA). Slides were visualized and imaged using a Nikon 80i microscope with a photometrics camera or a Nikon Eclipse TiE attached to an A1R-SHR confocal, with an A1-DU4 detector, and LU4 laser launch.

For quantification of cell proliferation, Ki67 labeled sections were counterstained with hematoxylin to visualize nucleus. Images were acquired from 4 random fields from each mouse section (total 4 mice each group) and total number of nuclei and Ki67 positive cells were counted in a blinded fashion. Data from each field was averaged and presented as percent Ki67 positive cells.

Cilia length quantification: Tissue sections immunostained for acetylated α-tubulin and DBA were imaged at multiple planes, which were then merged to obtain full-length cilia. Cilia images were converted to black and white and cilia lengths were quantified using ImageJ

### Western blots

Following treatments, cells were washed with PBS three times and lysed. Tissues (fresh or frozen) were chopped in pieces and homogenized using a Dounce homogenizer. For both cells and tissues, RIPA lysis buffer (50 mM Tris HCl pH7.5, 137 mM NaCl,1% IGEPAL, 2 mM EDTA) with protease inhibitors (Protease inhibitor cocktail, Fisher Scientific, Pittsburg, PA) was used (25). Protein concentration was measured using BCA protein assay (Bio-Rad, Hercules, CA). Whole cell lysates (50 to 100μg) were solubilized in 6X Laemmli sample buffer and heated to 95 °C for ten minutes and electrophoresed on 15% or 10% polyacrylamide gels. Proteins were transferred to PVDF membranes. Ponceau staining was performed for each blot to determine protein transfer and imaged. Ponceau S represented total protein and was used to normalize the gels for protein loading (26). The immunoblots were blocked in 5% nonfat dry milk in PBST (PBS containing 0.1% Tween 20) for 1 hour at room temperature and then followed by PBS washes; the blots were incubated with appropriate dilutions of primary antibodies overnight. The blots were then washed and incubated with secondary antibodies (1:10,000 dilution in blocking solution) for 1 hour at room temperature. After subsequent washes in PBST, bound antibody was detected by chemiluminescence (Western Lightning Plus ECL, Perkin Elmer). Bands produced in the results were quantified using imageJ and normalized with ponceau S staining to confirm equal loading (26). Data was presented as relative intensity.

### Statistics

Data are expressed as mean ± SE. Statistical significance was measured by Student’s unpaired T test for comparison between control and PKD groups. One-way ANOVA was performed to compare more than two groups followed by Tukey HSD test. A *P* value of <0.05 was considered statistically significant.

## Results

### Quinomycin A treatment reduces cyst growth from ADPKD cells grown in 3D collagen gels

Studies have shown that quinomycin A (fig: 1A) is an anticancer agent which intercalates double stranded DNA. As low as 5nM quinomycin A was able to induce cell cycle arrest in pancreatic cancer cells (14). We first determined the effects of Quinomycin A on kidney cell viability. Normal human kidney (NHK) cells and ADPKD cells obtained from cortical cysts of ADPKD patients were treated with increasing concentrations of Quinomycin A starting from 1 to 10nM (fig:1B). Treatments were continued for 4 consecutive days. Both NHK and ADPKD cells tolerated Quinomycin well at 1nM. At 2nM, while NHK cells maintained their viability, ADPKD cells lost approximately 25% viability. By 10nM, NHK cells still maintained 80% viability, whereas ADPKD lost 80% viability. The dose response curve showed that ADPKD cells are more responsive to Quinomycin A treatments. We next performed 3D cyst assays using primary ADPKD renal epithelial cells (22) to examine the potential of Quinomycin A on cysts. ADPKD cells were induced to form cysts in a collagen gel in the presence of the cAMP agonist, FSK, and growth factor, EGF. After cyst formation, Quinomycin A at desired concentrations or vehicle was used for 4 consecutive days and the effects of Quinomycin A on cyst progression were evaluated. As shown in figures 1C and 1D, Quinomycin A treatment reduced size of *in vitro* cysts, relative to vehicle treatment. Interestingly, while treatment with 2nm Quinomycin A resulted in only a 25% viability loss of ADPKD cells grown in 2D (figure 1B), the same treatment of Quinomycin A obliterated cysts when cells were grown in 3D (figures 1C and 1D). These data suggest that cysts may intrinsically be more dose-sensitive than cells grown in 2D.

**Figure 1.**
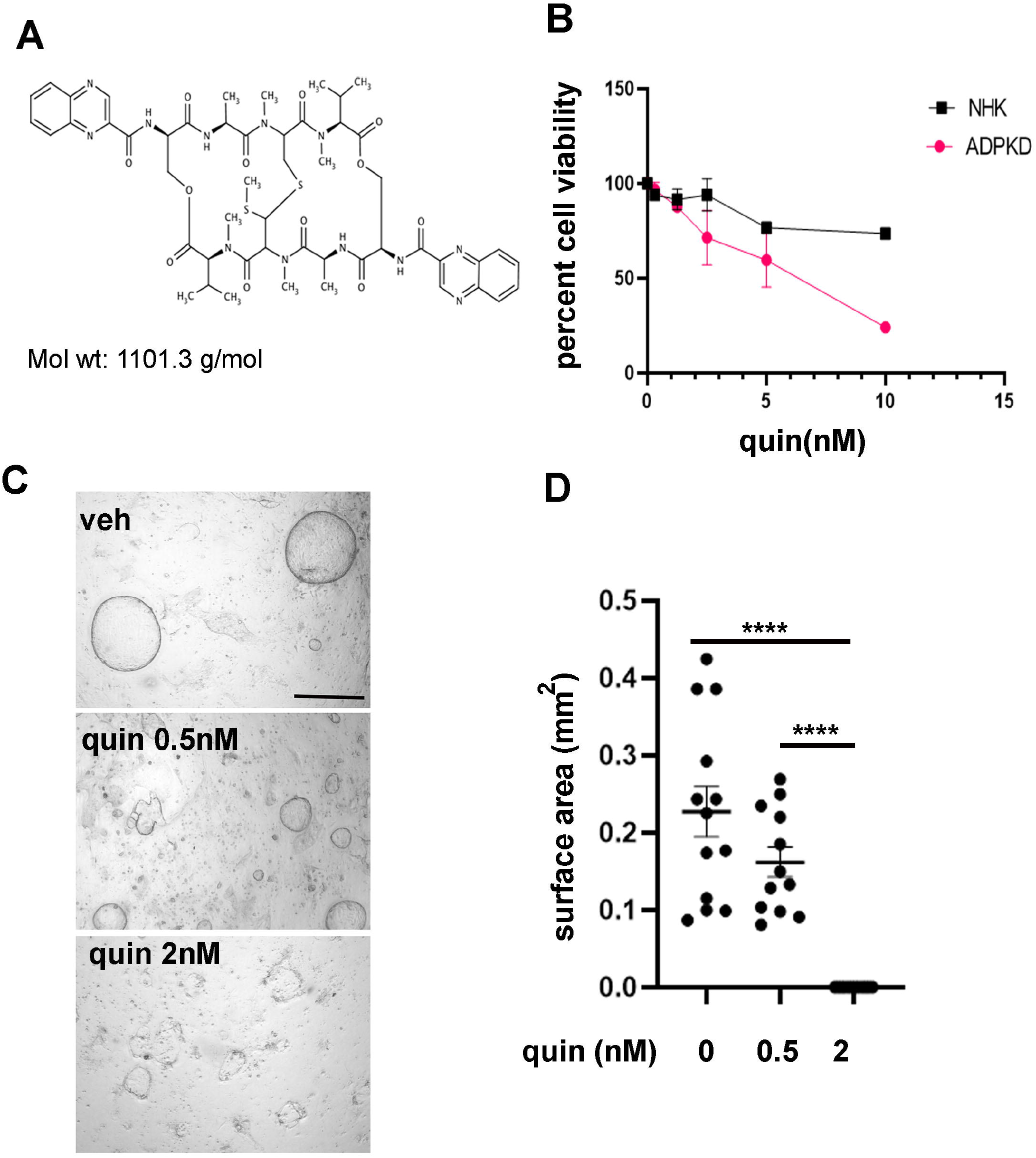
Primary ADPKD cells are sensitive to Quinomycin A treatments: **(A)** Chemical structure of Quinomycin A (C_51_H_64_N_12_O_12_S_2_) with molecular weight. **(B)** Primary cells from normal human kidney (NHK) and ADPKD (n=3 to 5) at each data point were cultured in presence of Quinomycin A for 4 days followed by viability assays. Data presented as percent viability. **(C)** Primary ADPKD cells were grown to form cysts on a 3-dimensional system using collagen matrix in the presence of forskolin and EGF for 3-5 days followed by treatment with vehicle or Quinomycin A (quin) for 4 additional days. Cysts were fixed, imaged and measured. **(D)** Cyst size represented as surface area from an ADPKD patient (six or more replicates/treatment). Each point represents a replicate. Statistical significance was determined using one-way ANOVA followed by Tukey’s HSD test. Data presented as mean +-SE. (*p<0.05), **(p<0.01). Scale bar: 0.57mm.

### Quinomycin A inhibits cyst progression in an ADPKD mouse model

We next examined the effects of Quinomycin A *in vivo* in a mouse model of ADPKD. We generated *PkdP^RC/RC^/Pkd2^+/-^* mice (hereafter called PKD mice) by breeding the *Pkd1^RC/RC^* mice with *Pkd2^+/-^* mice (both in C57BL/6J background) kindly provided by Drs. Peter Harris and Steven Somlo, respectively (15, 16). These mice show moderate cystic disease progression such that cysts are present at birth and continue to moderately progress until 6 weeks (unpublished). In mice, Quinomycin A concentration of 10μg/kg body weight intraperitoneal injections for up to 200 days did not result in any adverse effects (27). Based on these studies, PKD and littermate wildtype (WT) control mice were weaned at postnatal (P) day 21 and intraperitoneal injections of Quinomycin A (10μg/kg body weight) were administered from P22-P49. We did not notice any adverse events and mice remained active throughout the duration of the study. Mice were euthanized at P50, and tissues and blood samples were collected (fig. 2A). Kidneys of vehicle- and Quinomycin A-treated WT mice appeared similar. In contrast, a significant reduction in kidney size (fig. 2B) was observed in Quinomycin A-treated PKD kidneys compared to vehicle-treated PKD kidneys. As expected, the kidney size correlated with kidney-to-body weight ratios (fig. 2C), showing a significant reduction (p<0.05) in Quinomycin A-treated PKD mice relative to vehicle-treated PKD mice. This correlated with a significant reduction in cystic area in the Quinomycin A-treated PKD group in comparison to the vehicle treated PKD group (fig. 2D). Renal function, as assessed by blood urea nitrogen (BUN) levels, was not statistically different between groups, however Quinomycin A treatments showed a decreasing BUN trend in PKD mice (fig.2E). These data indicate a protective effect of Quinomycin A on PKD progression.

**Figure 2.**
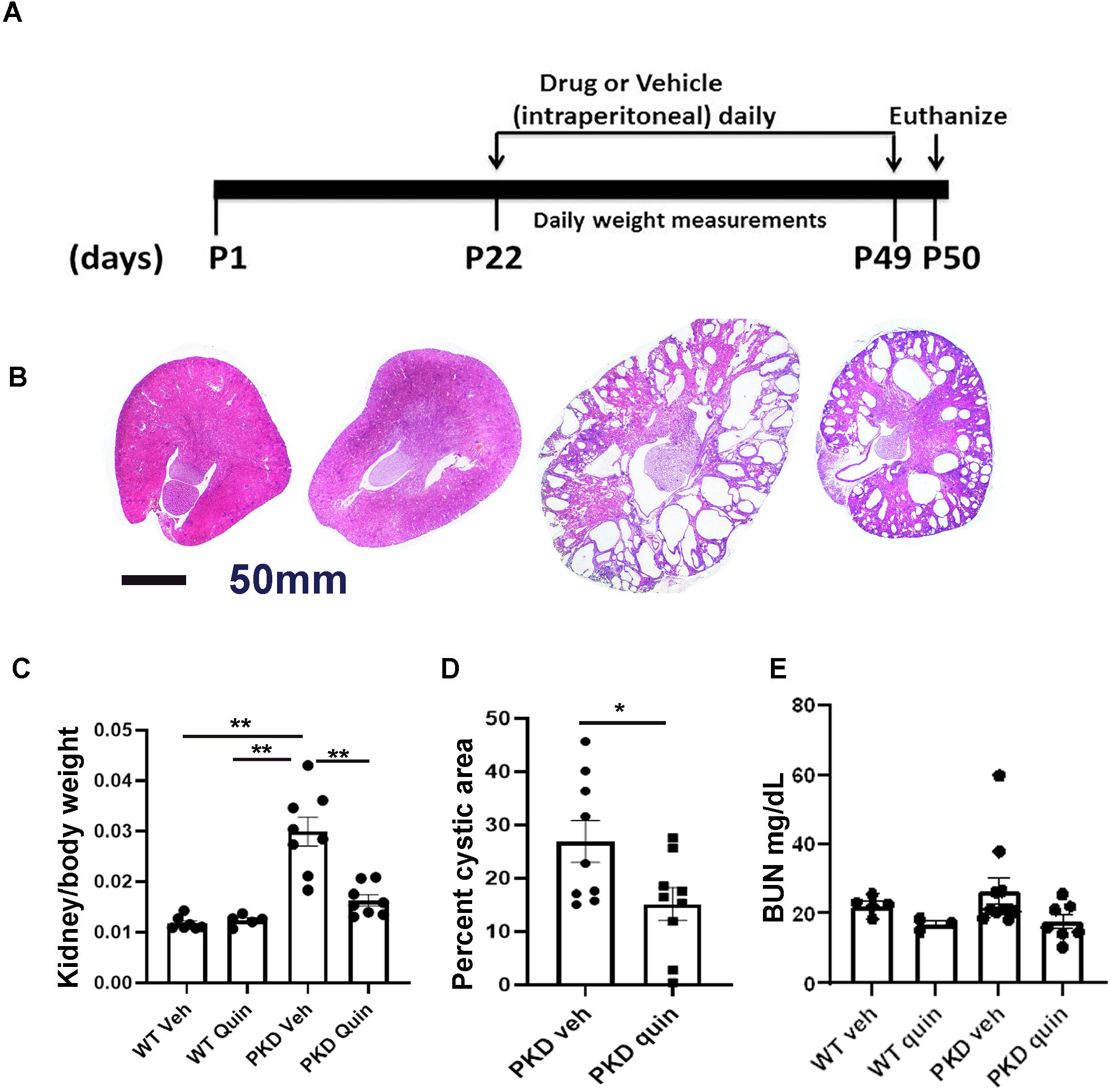
Quinomycin A reduces disease progression in a mouse model of ADPKD: **(A)** Experimental timeline for Quinomycin A treatments. PKD or WT mice (postnatal day 22) were intraperitoneally injected with vehicle (veh) or Quinomycin A (10 mg/kg body weight) for 27 consecutive days. Mice were euthanized and samples were collected. **(B)** H&E staining of kidney sections. Representative images of each treatment group are shown. **(C)** Total kidney to body weight ratio of vehicle and Quinomycin A treated PKD and WT mice **(D)** Cystic index of PKD kidneys measured as percent cystic area. **(E)** Serum urea nitrogen values measured as mg/dL. Each dot represents a mouse in C through E. Data presented as mean +_SE (n=4-8 per group). Statistical significance was determined using unpaired student’s T-test (D) or one-way ANOVA followed by Tukey’s HSD test (*p<0.05), (**p<0.01). Scale bar: 1mm.

### Quinomycin A reduces cell proliferation, fibrosis in PKD kidneys

The intercalating functions of Quinomycin A affects dividing cells. We asked whether Quinomycin A can counter cell proliferation, a hallmark of PKD. Kidneys from control and PKD mice were labelled for Ki67, a proliferation marker. Vehicle-treated PKD mice exhibited approximately 9% positive cells for Ki67/ kidney whereas Quinomycin A-treated PKD mice showed a significant reduction with only ~5% Ki67 positive cells on immunolabeling (fig: A and B). PKD is also characterized by renal myofibrogenic activity which contributes to fibrosis. We immunolabelled the sections for presence of myofibroblast marker, alpha smooth muscle actin (α-SMA) which is normally expressed in smooth muscle cells (SMCs) surrounding blood vessels (V) as seen in figure 3C. PKD kidneys were highly positive for α-SMA labelling, not restricted to SMCs, but also present around cysts (fig:3C). In contrast to vehicle-treated PKD kidneys, Quinomycin A treatment in PKD mice reduced α-SMA labeling. Western blot analyses of kidney lysates from Quinomycin A- and vehicle-treated WT and PKD mice (fig: 3D) showed low expression of α-SMA in Quinomycin A and vehicle treated WT kidneys, but a significantly (P<0.001) marked increase of α-SMA in PKD (veh) kidneys. In contrast, Quinomycin A treatment significantly decreased α-SMA expression which were comparable to control. These data show that Quinomycin A partially rescues PKD progression by decreasing cell proliferation and myofibrotic activity

**Figure 3.**
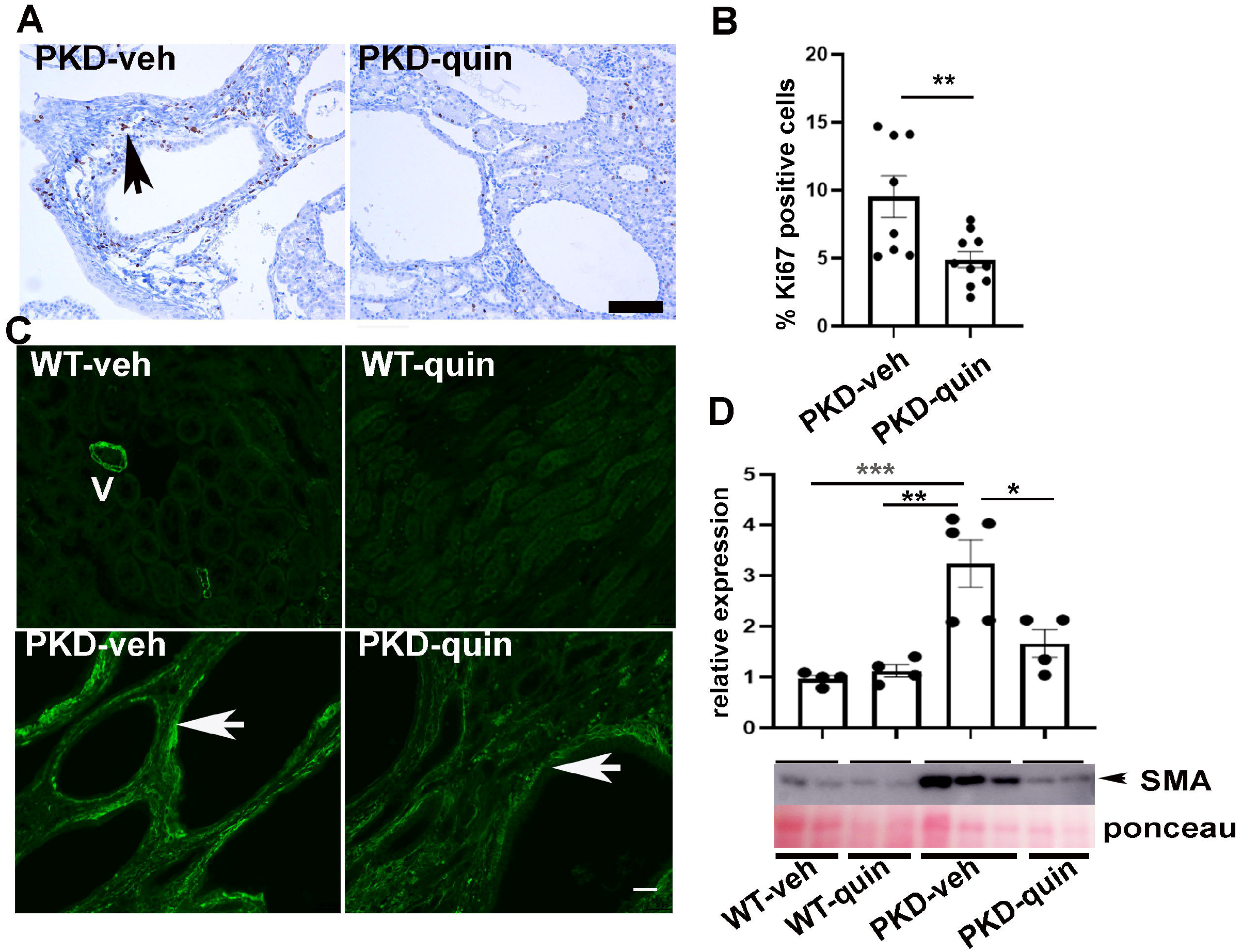
Quinomycin A slows down cell proliferation and myofibrosis in PKD: **(A)** Ki67 staining was performed to via Immunohistochemistry (IHC) to asses cell proliferation in Quinomycin A and vehicle treated WT and PKD kidneys. Arrow points to highly proliferative area on a vehicle (veh) treated PKD mouse kidney section. These proliferating areas were reduced in Quinomycin A treated PKD mice. Hematoxylin staining shows blue nuclei and Ki67 positive nuclei are shown in dark brown. **(B)** quantification of Ki67 staining expressed as percent Ki67 positive cells from at least 7 to 10 images randomly taken from 4 mice in each group. Unpaired Student T test used for statistical analysis. Data presented as mean +-SE. (*p<0.05), ((**p<0.01), (***p<0.001). Scale bar: 100μm.

### Quinomycin treatment inhibits Notch signaling in ADPKD mice

We have previously reported that the Notch signaling pathway is activated in renal cyst-lining epithelial cells and interstitial cells of mouse models and in human ADPKD renal tissue and cells (9). Since Quinomycin A can target the Notch signaling pathway (14), we examined whether Quinomycin A can affect Notch signaling pathway in our model. IHC for Notch3 revealed reduced Notch3 labeling, especially in the cystic epithelial cells (fig 4A, arrows). Consistent with our previous report, in Western blots of total kidney lysates although a decreasing trend in Notch3 IC was observed, it did not reach significance (fig:3B). However, other members of the Notch pathway, RBPjk and HeyL were significantly elevated in PKD lysates ( fig:4C and 4D), and showed a reduction in kidneys treated with Quinomycin A. Our data show that Quinomycin A effectively targets the Notch signaling and protects kidneys from further injury.

**Figure 4.**
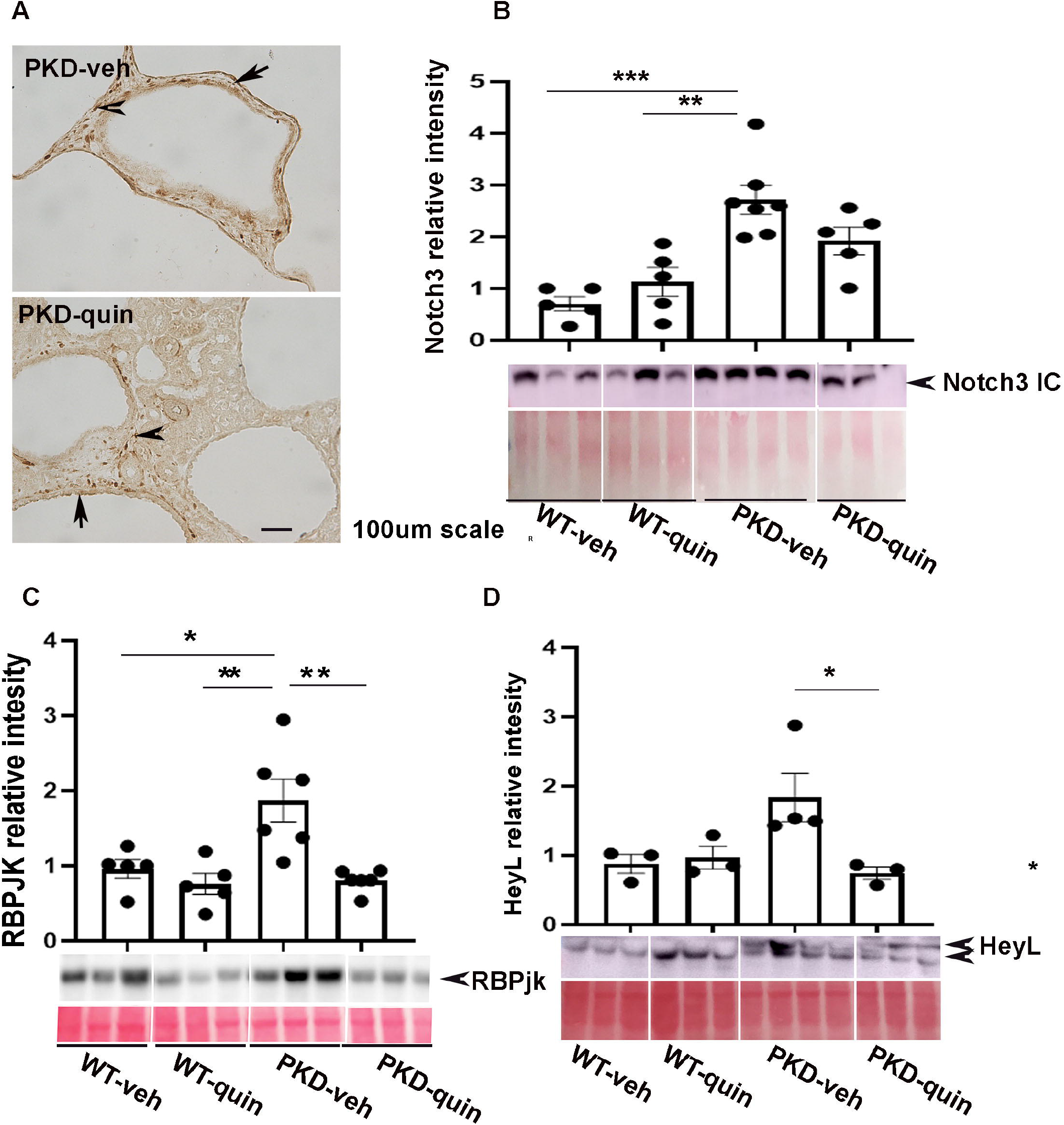
Quinomycin A targets the Notch signaling pathway in PKD: **(A)** Immunohistochemistry (IHC) for Notch3 (N3 IC). **(B)** Western blot (WB) analysis for Notch3 IC and cumulative quantitative data, **(C and D)** Western blot analysis for RBPjk and HeyL (Notch targets) showing increased expression in kidney lysates of vehicle treated PKD mice and Quinomycin A treatments reducing this expression. Protein blots were quantified using Ponceau S staining for total protein normalization. Data presented as relative change. Each dot represents individual mouse. Data presented as mean +-SE. Statistical significance was determined using One-way ANOVA followed by Tukey’s HSD test (*p<0.05), (**p<0.01), (***p<0.001). Scale bar: 100μm.

### Quinomycin normalizes cilia length in an ADPKD mouse model

Studies have shown that cilia are lengthened in kidney epithelial cells of PKD mouse models, which may reflect defects in ciliary homeostasis (15, 28–30). Thus, we assessed cilia length and the effect of Quinomycin A on cilia length in mouse kidneys. Kidney sections were immunolabeled for the presence of acetylated α-tubulin which marks cilia together with DBA lectin which stains collecting ducts, where cysts predominantly develop in ADPKD. Immunofluorescence labelling revealed that cilia were lengthened in vehicle-treated PKD kidneys compared to the vehicle-treated WT kidneys (figs:5A and 5B), Quinomycin A treatments shortened cilia lengths in PKD kidneys (fig:5A), Cilium are pointed by arrows and are represented in magnified insets under each image; (fig 5A). Quantification of cilia length within kidneys further confirmed highly significant (P<0.0001) cilia length reduction following Quinomycin A treatments in PKD mice, indicating that Quinomycin A restores cilia lengths in a PKD mouse model to those of wild type mice (fig:5B).

**Figure 5.**
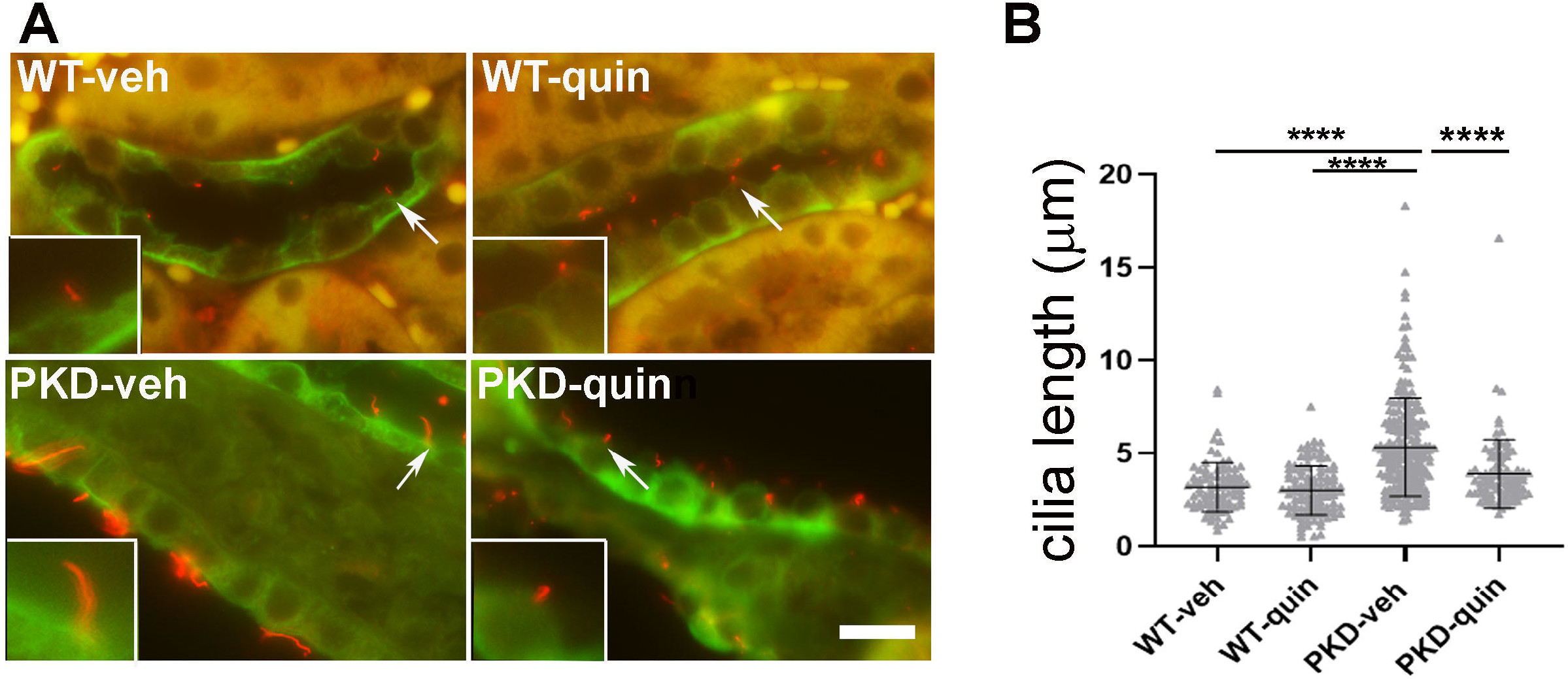
Quinomycin A normalizes cilia length in PKD kidneys: **(A)** Double immunostaining of primary cilia marker protein, acetylated tubulin (red) together with *Dolichus biflorus agglutinin* (DBA)(green). Arrows represent cilium which are magnified and shown in insets. (B) Cilia length was quantified from n=4 mice per group. Each data point represents a cilium. Data presented as mean +-SD. Statistical significance was determined using One-way ANOVA followed by Tukey’s HSD test (p<0.0001). Scale bar: 10μM

## Discussion

In this study, we show that Quinomycin A effectively reduces cyst size in primary epithelial cells obtained from cysts of ADPKD patients. We also show that normal human kidney epithelial cells are not as sensitive to Quinomycin A as ADPKD cells, indicating Quinomycin A can spare normal epithelial cells, while targeting the cystic cells. In support of our study, *in vitro* studies have been performed with as low as 5nM Quinomycin A in several cancerous cell lines (31, 32). In pancreatic cell lines this concentration was effective for inhibition of cancer cell proliferation but did not affect normal human pancreatic ductal epithelial cells even at a concentration of 50nM, supporting our observation that normal cells are not as responsive to Quinomycin A. The data are further supported by the 3D *in vitro* cyst assays in which ADPKD cells grown in cysts were responsive to even lower doses of Quinomycin than ADPKD cells grown in 2D. This effect could be attributed to dependence of cyst formation on cyclic AMP. While cell viability is tested in cells grown in media without addition of other signaling factors, initiation of cyst formation *in vitro* (3D) takes place in response to forskolin, an agonist of cyclic AMP, an important mediator of cyst progression (22, 33). Thus, it is possible that Quinomycin A is sensitive to cAMP. Further studies will be required to support this observation.

Quinomycin A is a member of quinoxaline family of antibiotics previously identified as an anti-cancer drug that binds strongly to double stranded DNA (12). Due to its strong DNA chelator property, Quinomycin A can be harmful in high doses and prior clinical trials have documented multiple toxicities because of which the drug development was halted. These toxicities were attributed to the higher doses 180 mg/m^2^ of Quinomycin A (34–40). A dose of 60-120mg/m^2^ was found to have no toxicity (41). Recent studies in mice indicate that low doses (10μg/kg body weight) of Quinomycin A are not toxic to normal cell hematopoiesis and selectively targets cancerous cells in acute myelocytic leukemia and bladder cancer (14, 27). To further combat the negative effects of Quinomycin A, a recent study showed that actively targeted nano-delivery of Quinomycin A can efficiently target cancer cells to induce autophagic cell death instead of apoptosis or necrotic cell death (42). Based on these studies it is possible that alternative doses and targeted nano-delivery of Quinomycin A can have better therapeutic potential in clinical trials.

In addition to targeting Notch pathway, Quinomycin A has been shown to target HIF-1α. HIF-1α expression is increased in cystic epithelia and was found to be associated with increased cyst growth by activating fluid secretion (43). We have previously shown Notch3 activation in cyst lining, cells as well as in interstitial cells, in PKD (9). It is possible that cross talk between HIF and Notch signaling exists in cystic tissues as documented previously where HIF-1α binds to cleaved Notch which results in stabilization and activation of Notch signaling (44, 45). In another study, chronic hypoxia was shown to activate Notch signaling in prostate cancer. Although we were not successful in detecting HIF-1α in the kidney nuclear extracts of the PKD mice via Western blots or immunohistochemistry, Quinomycin A reduced the Notch pathway downstream members, RBPjk and HeyL showing that Notch pathway is a target of Quinomycin.

Elongation of cilia has been observed on tubular epithelial cells in PKD (28, 29). The primary cilium is an antenna-like mechano- and chemosensor that detects fluid flow and transduces extracellular signals into the cells in order to maintain renal function and nephron structure (46, 47). Verghese et al. suggested that cilia elongation can be a critical factor in conferring kidney disease resistance (48, 49). Moreover, Notch activation leads to lengthening of cilia in developing neural tube in mouse embryos and in NIH3T3 cells (50). Thus, it is possible that Quinomycin A reduces Notch signaling, which in turn, results in shortening of cilia in PKD kidneys in our mouse model. This hypothesis will need further evaluation.

Taken together, we show that Quinomycin A inhibits the Notch signaling pathway, reduces cilia length and confers protection against cyst progression in ADPKD by reducing cellular proliferation and fibrosis. To our knowledge, this is the first evidence where Quinomycin A has been shown to alter cilia length. The study adds QuinomycinA to the list of drugs that may prove beneficial to PKD patients in clinics.

## Author Contributions

M.S conceived and designed the study. M.S, P.S.R, M.M.T, and B.M conducted experiments, P.V.T and D.P.W. provided reagents. M.S., D.S., P.V.T., D.P.W. and J.P.C analyzed data and interpreted results of experiments, M.S prepared figures; M.S drafted manuscript; M.S, P.V.T, D.S, D.P.W and J.P.C edited and revised manuscript; M.S. approved the final version of manuscript. All authors read and approved the final manuscript.

## Acknowledgements

We thank Dr. Rajni Vaid Puri, Anh Dao Do, Jessica Idowu, Wei Wang and Johnny Dinh Phan for technical assistance. Gail Reif, Yan Zhang and Emily Daniel for help with NHK and ADPKD cell acquisition. Jing Huang in histology core for sectioning of the kidneys. This work was supported by the K-INBRE summer scholarship award (P20 GM103418) to PR, PKD Biomaterials and Biomarkers Repository Core in the Kansas PKD Research and Translational Core Center (NIH P30DK106912 to JPC), R01DK103033 to PVT, and R01DK108433 to MS.

## References

1. Gallagher AR, Germino GG, and Somlo S. Molecular advances in autosomal dominant polycystic kidney disease. Adv Chronic Kidney Dis. 2010;17(2): 118–30.

2. Harris PC, and Torres VE. Polycystic kidney disease. Annu Rev Med. 2009;60:321–37.

3. Nauli SM, Alenghat FJ, Luo Y, Williams E, Vassilev P, Li X, et al. Polycystins 1 and 2 mediate mechanosensation in the primary cilium of kidney cells. Nature genetics. 2003;33(2):129–37.

4. Torres VE, Chapman AB, Devuyst O, Gansevoort RT, Grantham JJ, Higashihara E, et al. Tolvaptan in patients with autosomal dominant polycystic kidney disease. N Engl J Med. 2012;367(25):2704–18.

5. Reif GA, Yamaguchi T, Nivens E, Fujiki H, Pinto CS, and Wallace DP. Tolvaptan inhibits ERK-dependent cell proliferation, Cl-secretion, and in vitro cyst growth of human ADPKD cells stimulated by vasopressin. Am J Physiol Renal Physiol. 2011;301(5):F1005–13.

6. Sorohan BM, Ismail G, Andronesi A, Micu G, Obrişcă B, Jurubiţă R, et al. A single-arm pilot study of metformin in patients with autosomal dominant polycystic kidney disease. BMC nephrology. 2019;20(1):276.

7. Zhou X, Fan LX, Sweeney WE Jr, Denu JM, Avner ED, and Li X. Sirtuin 1 inhibition delays cyst formation in autosomal-dominant polycystic kidney disease. J Clin Invest. 2013;123(7):3084–98.

8. Sweeney WE, Frost P, and Avner ED. Tesevatinib ameliorates progression of polycystic kidney disease in rodent models of autosomal recessive polycystic kidney disease. World journal of nephrology. 2017;6(4):188–200.

9. Idowu J, Home T, Patel N, Magenheimer B, Tran PV, Maser RL, et al. Aberrant Regulation of Notch3 Signaling Pathway in Polycystic Kidney Disease. Scientific Reports. 2018;8(1):3340.

10. Ran Y, Hossain F, Pannuti A, Lessard CB, Ladd GZ, Jung JI, et al. γ-Secretase inhibitors in cancer clinical trials are pharmacologically and functionally distinct. EMBO molecular medicine. 2017;9(7):950–66.

11. Foster BJ, Clagett-Carr K, Shoemaker DD, Suffness M, Plowman J, Trissel LA, et al. Echinomycin: the first bifunctional intercalating agent in clinical trials. Investigational new drugs. 1985;3(4):403–10.

12. Waring MJ, and Wakelin LP. Echinomycin: a bifunctional intercalating antibiotic. Nature. 1974;252(5485):653–7.

13. Wang Y, Liu Y, Tang F, Bernot KM, Schore R, Marcucci G, et al. Echinomycin protects mice against relapsed acute myeloid leukemia without adverse effect on hematopoietic stem cells. Blood. 2014;124(7):1127–35.

14. Ponnurangam S, Dandawate PR, Dhar A, Tawfik OW, Parab RR, Mishra PD, et al. Quinomycin A targets Notch signaling pathway in pancreatic cancer stem cells. Oncotarget. 2016;7(3):3217–32.

15. Hopp K, Ward CJ, Hommerding CJ, Nasr SH, Tuan HF, Gainullin VG, et al. Functional polycystin-1 dosage governs autosomal dominant polycystic kidney disease severity. J Clin Invest. 2012;122(11):4257–73.

16. Wu G, Markowitz GS, Li L, D’Agati VD, Factor SM, Geng L, et al. Cardiac defects and renal failure in mice with targeted mutations in Pkd2. Nat Genet. 2000;24(1):75–8.

17. Sharma M, Callen S, Zhang D, Singhal PC, Vanden Heuvel GB, and Buch S. Activation of Notch signaling pathway in HIV-associated nephropathy. AIDS. 2010;24C(14):2161–70.

18. Reif GA, Yamaguchi T, Nivens E, Fujiki H, Pinto CS, and Wallace DP. Tolvaptan inhibits ERK-dependent cell proliferation, Cl^−^ secretion, and in vitro cyst growth of human ADPKD cells stimulated by vasopressin. Am J Physiol Renal Physiol. 2011;301(5):F1005–13.

19. Yamaguchi T, Nagao S, Wallace DP, Belibi FA, Cowley BD, Pelling JC, et al. Cyclic AMP activates B-Raf and ERK in cyst epithelial cells from autosomal-dominant polycystic kidneys. Kidney Int. 2003;63(6):1983–94.

20. Reif GA, and Wallace DP. ADPKD cell proliferation and Cl(-)-dependent fluid secretion. Methods Cell Biol. 2019;153:69–92.

21. Reif GA, Yamaguchi T, Nivens E, Fujiki H, Pinto CS, and Wallace DP. Tolvaptan inhibits ERK-dependent cell proliferation, Cl^-^ secretion, and in vitro cyst growth of human ADPKD cells stimulated by vasopressin. Am J Physiol Renal Physiol. 2011;301:F1005–13.

22. Sharma M, Reif GA, and Wallace DP. In vitro cyst formation of ADPKD cells. Methods Cell Biol. 2019;153:93–111.

23. Wallace DP, Grantham JJ, and Sullivan LP. Chloride and fluid secretion by cultured human polycystic kidney cells. Kidney Int. 1996;50(4):1327–36.

24. Sharma M, Magenheimer LK, Home T, Tamano KN, Singhal PC, Hyink DP, et al. Inhibition of Notch pathway attenuates the progression of human immunodeficiency virus-associated nephropathy. Am J Physiol Renal Physiol 2013;304(8):F1127–36.

25. Idowu J, Home T, Patel N, Magenheimer B, Tran PV, Maser RL, et al. Aberrant regulation of Notch3 signaling pathway in polycystic kidney disease. Sci Rep. 2018;DOI: 10.1038/s41598-018-21132-3.

26. Sander H, Wallace S, Plouse R, Tiwari S, and Gomes AV. Ponceau S waste: Ponceau S staining for total protein normalization. Anal Biochem. 2019;575:44–53.

27. Wang Y, Liu Y, Tang F, Bernot KM, Schore R, Marcucci G, et al. Echinomycin protects mice against relapsed acute myeloid leukemia without adverse effect on hematopoietic stem cells. Blood. 2014;124(7).

28. Husson H, Moreno S, Smith LA, Smith MM, Russo RJ, Pitstick R, et al. Reduction of ciliary length through pharmacologic or genetic inhibition of CDK5 attenuates polycystic kidney disease in a model of nephronophthisis. Human molecular genetics. 2016;25(11):2245–55.

29. Gerakopoulos V, Ngo P, and Tsiokas L. Loss of polycystins suppresses deciliation via the activation of the centrosomal integrity pathway. Life science alliance. 2020;3(9).

30. Shao L, El-Jouni W, Kong F, Ramesh J, Kumar RS, Shen X, et al. Genetic reduction of cilium-length by targeting intraflagellar transport 88 protein impedes kidney and liver cysts formation in mouse models of autosomal polycystic kidney disease. Kidney international. 2020.

31. Yonekura S, Itoh M, Okuhashi Y, Takahashi Y, Ono A, Nara N, et al. Effects of the HIF1 inhibitor, echinomycin, on growth and NOTCH signalling in leukaemia cells. Anticancer Res. 2013;33(8):3099–103.

32. Kong D, Park EJ, Stephen AG, Calvani M, Cardellina JH, Monks A, et al. Echinomycin, a smallmolecule inhibitor of hypoxia-inducible factor-1 DNA-binding activity. Cancer Res. 2005;65(19):9047–55.

33. Putnam WC, Swenson SM, Reif GA, Wallace DP, Helmkamp GM, Jr., and Grantham JJ. Identification of a forskolin-like molecule in human renal cysts. Journal of the American Society of Nephrology : JASN. 2007;18(3):934–43.

34. Chang AY, Kim K, Boucher H, Bonomi P, Stewart JA, Karp DD, et al. A randomized phase II trial of echinomycin, trimetrexate, and cisplatin plus etoposide in patients with metastatic nonsmall cell lung carcinoma: an Eastern Cooperative Oncology Group Study (E1587). Cancer. 1998;82(2):292–30.

35. Gradishar WJ, Vogelzang NJ, Kilton LJ, Leibach SJ, Rademaker AW, French S, et al. A phase II clinical trial of echinomycin in metastatic soft tissue sarcoma. An Illinois Cancer Center Study. Invest New Drugs. 1995;13(2):171–4.

36. Wadler S, Tenteromano L, Cazenave L, Sparano JA, Greenwald ES, Rozenblit A, et al. Phase II trial of echinomycin in patients with advanced or recurrent colorectal cancer. Cancer Chemother Pharmacol. 1994;34(3):266–9.

37. Shevrin DH, Lad TE, Guinan P, Kilton LJ, Greenburg A, Johnson P, et al. Phase II trial of echinomycin in advanced hormone-resistant prostate cancer. An Illinois Cancer Council study. Invest New Drugs. 1994;12(1):65–6.

38. Chang AY, Tu ZN, Bryan GT, Kirkwood JM, Oken MM, and Trump DL. Phase II study of echinomycin in the treatment of renal cell carcinoma ECOG study E2885. Invest New Drugs. 1994;12(2):151–3.

39. Muss HB, Blessing JA, and DuBeshter B. Echinomycin in recurrent and metastatic endometrial carcinoma. A phase II trial of the Gynecologic Oncology Group. Am J Clin Oncol. 1993;16(6):492–3.

40. Marshall ME, Wolf MK, Crawford ED, Taylor S, Blumenstein B, Flanigan R, et al. Phase II trial of echinomycin for the treatment of advanced renal cell carcinoma. A Southwest Oncology Group study. Invest New Drugs. 1993;11(2-3):207–9.

41. Kuhn JG, Von Hoff DD, Hersh M, Melink T, Clark GM, Weiss GR, et al. Phase I trial of echinomycin (NSC 526417), a bifunctional intercalating agent, administered by 24-hour continuous infusion. Eur J Cancer Clin Oncol. 1989;25(5):797–803.

42. Thomas A, Samykutty A, Gomez-Gutierrez JG, Yin W, Egger ME, McNally M, et al. Actively Targeted Nanodelivery of Echinomycin Induces Autophagy-Mediated Death in Chemoresistant Pancreatic Cancer In Vivo. Cancers. 2020;12(8).

43. Buchholz B, Schley G, Faria D, Kroening S, Willam C, Schreiber R, et al. Hypoxia-inducible factor-1α causes renal cyst expansion through calcium-activated chloride secretion. J Am Soc Nephrol. 2014;25(3):465–74.

44. Gustafsson MV, Zheng X, Pereira T, Gradin K, Jin S, Lundkvist J, et al. Hypoxia requires notch signaling to maintain the undifferentiated cell state. Dev Cell. 2005;9(5):617–28.

45. Wang Y, Liu Y, Malek SN, Zheng P, and Liu Y. Targeting HIF1α eliminates cancer stem cells in hematological malignancies. Cell Stem Cell. 2011;8(4):399–411.

46. Praetorius HA, and Spring KR. A physiological view of the primary cilium. Annual review of physiology. 2005;67:515–29.

47. Park KM. Can Tissue Cilia Lengths and Urine Cilia Proteins Be Markers of Kidney Diseases? Chonnam medical journal. 2018;54(2):83–9.

48. Verghese E, Zhuang J, Saiti D, Ricardo SD, and Deane JA. In vitro investigation of renal epithelial injury suggests that primary cilium length is regulated by hypoxia-inducible mechanisms. Cell Biol Int. 2011;35(9):909–13.

49. Overgaard CE, Sanzone KM, Spiczka KS, Sheff DR, Sandra A, and Yeaman C. Deciliation is associated with dramatic remodeling of epithelial cell junctions and surface domains. Mol Biol Cell. 2009;20(1):102–13.

50. Stasiulewicz M, Gray SD, Mastromina I, Silva JC, Björklund M, Seymour PA, et al. A conserved role for Notch signaling in priming the cellular response to Shh through ciliary localisation of the key Shh transducer Smo. Development (Cambridge, England). 2015;142(13):2291–303.

